# Anti-tumor efficacy of an MMAE conjugated antibody targeting cell surface TACE/ADAM17-cleaved Amphiregulin in breast cancer

**DOI:** 10.1101/2021.07.07.451518

**Authors:** Kristopher A. Lofgren, Sreeja Sreekumar, E. Charles Jenkins, Kyle J. Ernzen, Paraic A. Kenny

**Author notes:** Equal Contribution (Kristopher A. Lofgren, Sreeja Sreekumar). Corresponding Author: Paraic A. Kenny.

## Abstract

The Epidermal Growth Factor Receptor ligand, Amphiregulin, is a key proliferative effector of estrogen receptor signaling in breast cancer and also plays a role in other malignancies. Amphiregulin is a single-pass transmembrane protein proteolytically processed by TACE/ADAM17 to release the soluble EGFR ligand, leaving a residual transmembrane stalk that is subsequently internalized. Here, we report the development of an antibody drug conjugate, GMF-1A3-MMAE, targeting an AREG neo-epitope revealed following ADAM17-mediated cleavage. The antibody does not interact with uncleaved Amphiregulin, providing a novel means of targeting cells with high rates of Amphiregulin shedding. Using fluorescent dye conjugation, we demonstrated that the antibody is internalized by cancer cells in a manner dependent on the presence of cell surface cleaved Amphiregulin. Antibodies conjugated with monomethyl auristatin E (MMAE) were cytotoxic in vitro and induced rapid regression of established breast tumor xenografts in immunocompromised mice. We further demonstrate that these antibodies recognize the Amphiregulin neo-epitope in formalin fixed paraffin embedded tumor tissue, suggesting their utility as a companion diagnostic for patient selection.

## Introduction

Breast cancer is the second most frequently diagnosed cancer in women and is classified into clinical subtypes by the expression status of the estrogen and progesterone receptors (ER, PR) and the presence or absence of HER2 amplification. ER+ tumors comprise approximately 70% of all breast tumors (Waks and Winer, 2019). Patients with ER+ disease receive treatment with endocrine targeting therapies such as tamoxifen and aromatase inhibitors with significant initial success, however the emergence of ER independent or endocrine therapy resistant disease is problematic (Clarke et al., 2015). As such, the development of alternative therapeutics for endocrine-insensitive disease is of critical importance.

Amphiregulin (AREG) is a transmembrane protein which, following TACE/ADAM17-dependent cleavage (Gschwind et al., 2003), releases a soluble EGFR ligand domain which promotes proliferation of normal and malignant cells. Inhibition of ADAM17 with protease inhibitors prevents Amphiregulin shedding and effectively interrupts this EGFR activating signal (Kenny and Bissell, 2007). Amphiregulin is transcriptionally regulated by estrogen during normal mammary gland development (McBryan et al., 2007) and in breast cancer (Peterson et al., 2015) and is a required effector for estrogen’s proliferation signal in both settings. High Amphiregulin expression levels are correlated to ER-alpha expression (Kenny and Bissell, 2007; Peterson et al., 2015) and, when overexpressed experimentally, Amphiregulin promotes EGF signaling self-sufficiency (Willmarth and Ethier, 2006). While our primary focus has been on the role of AREG in breast cancer, we note that it is expressed in several other cancer types (Busser et al., 2011) so the potential utility of therapeutic antibody drug conjugates extends to other malignancies.

Antibody-drug conjugates (ADCs) consist of an antibody against a specifically targeted epitope that is used to deliver locally high concentrations of a toxic payload to a cell of interest. The payload can be toxins or chemicals that induce DNA damage, disrupt cytoskeletal integrity, or interfere with the process of DNA replication. The value of ADCs is the specific delivery to a tumor of drugs that may be too toxic for patients or have complicating side effects if given at a dose high enough for tumor eradication. At least 34 unique ADCs are in various stages of clinical trials in the United States for treating solid tumors, with only three being FDA approved for clinical use in breast cancer at the time of publication: two targeting HER2 (trastuzumab emtansine, trastuzumab deruxtecan) and sacituzumab govitecan, which targets Trop-2 in metastatic TNBC patients (Barroso-Sousa and Tolaney, 2021). ADC development in breast cancer has mostly been directed towards HER2+ disease, with relatively little attention given to ER+ disease due to a paucity of targets. Thus, the identification of novel epitopes to be targeted in the context of ER+ disease is of value.

Given the strong requirement for Amphiregulin expression and proteolytic cleavage in ER+ breast cancer (Peterson et al., 2015), we postulated that the residual cell-surface stalk resulting from AREG cleavage could serve as a neo-epitope for antibody drug conjugate development. Here, we describe the identification of antibodies that selectively recognize cleaved, cell-associated Amphiregulin and the evaluation of their utility as antibody-drug conjugates in preclinical breast cancer models.

## 2. Materials and Methods

### 2.1 Cell lines and culture conditions

Human breast cancer cell line MCF7 was obtained from American Type Culture Collection (Manassas, VA, USA). The cell line was cultured in high-glucose Dulbecco’s modified Eagle medium (DMEM) supplemented with 10% fetal bovine serum (Gibco, Life Technologies, Carlsbad, CA, USA) at 37°C in 5% CO2. The cell line was authenticated by short-tandem repeat profiling (Arizona Genetics Core, Tucson, Arizona, USA). The MCF7 fulvestrant-resistant subline (MCF7-F) was kindly provided by Dr. Kenneth P. Nephew and cultured as previously described (Fan et al., 2006).

### 2.2 Antibodies

Rabbit monoclonal IgG antibodies specific to cleaved AREG were identified in contracted screening (Oak Biosciences Inc., Sunnyvale, CA, USA) using a recombinant Rabbit naïve phage display library (scFv format, VK-linker-VH) screening platform after 4 rounds of panning and ELISA validation. For panning, phage were selected against a peptide modeling the ADAM17-cleaved cell-associated epitope of AREG (Levano and Kenny, 2012), represented by the N-terminal region of THSMIDSSLSKI, and counterselected against a peptide spanning this region (SMKTHSMIDSSLSKIAC), representing uncleaved AREG. Three candidate clones – GMF-1A3, GMF-3A3 and GMF-3E4 – were reformatted as IgG and selectivity for cleaved Amphiregulin was confirmed by ELISA. GMF-1A3, referred to hereafter as simply “1A3”, was the antibody chosen for further characterization in this study.

### 2.3 Immunohistochemistry

A breast disease spectrum tissue microarray was purchased from US Biomax Inc. (Derwood, MD, USA). The slides were deparaffinized in xylene and rehydrated by serial incubations in graded ethanol and then in water in a Histo-Tek® SL Slide Stainer (Sakura Finetek USA, Inc., Torrance, CA, USA). Antigen retrieval was performed in a steamer by boiling slides in a container of citrate buffer (pH 6.0) for 20 min which was then removed for 15 minutes of cooling on the benchtop. Slides were washed in 1x Wash buffer (Dako Agilent, Santa Clara, CA, USA) and endogenous peroxidase was quenched by incubating with Dako Dual Endogenous Enzyme Block (S2003) for 10 minutes. Slides were washed in 1x Wash buffer, blocked (5% rabbit/10% goat serum in PBS), and immunostained with goat anti-AREG antibody (15 μg/mL; AF262, R&D Systems) or rabbit anti-cleaved AREG 1A3 antibody (10 μg/mL) overnight at 4°C. Slides were washed four times in 1x Wash buffer, followed by incubation for 45 minutes at room temperature in 1:100 dilution of rabbit anti-goat immunoglobulins/HRP (Dako, P0160 or ready-to-use goat anti-rabbit HRP labelled polymer (Dako, K4003). The slides were washed twice in 1x Wash buffer and the color was developed with 3,3-diaminobenzidine tetrahydrochloride (DAB) substrate chromogen system (K3468, DAKO, Botany, Australia). Sections were washed with water and counterstained with hematoxylin, rinsed with water, dehydrated by serial ethanol washes to 100%, cleared, and mounted in Permount (Thermo Fisher Scientific). The staining intensity was assessed semi-quantitatively using a four-point scale (Negligible=0, Low=1, Medium=2, High=3) by two investigators working independently on blinded samples. Discordant scores were resolved by joint review.

### 2.4 Internalization and intracellular localization analysis of anti-cleaved-AREG 1A3 antibody

In order to study internalization, 1A3 was conjugated with pHrodo iFL Red dye using pHrodo iFL Red Microscale Protein Labeling Kit (P36014, Thermo Fisher Scientific). MCF7 cells were seeded in 12 mm Nunc Glass Bottom Dish, at a density of 1×10^5^ cells/dish. After allowing the cells to attach overnight, the cells were incubated for 60 min with 1 mg/ml Hoechst dye for nuclei staining and LysoTracker® Green DND-26 (Thermo Fisher Scientific) for lysosomal labelling. The cells were then incubated with 1A3-pHrodo (2 μg/ml) for 20 min at 37°C. After washing the wells with PBS, the cells were supplemented with 1x live cell imaging solution (Molecular probes, Thermo Fisher Scientific). Fluorescence images were acquired at 30 min intervals for 2.5 h with the Olympus 1X71 inverted fluorescence microscope (Olympus, Shinjuku, Tokyo, Japan). TAPI-2 (20 μM)-pretreatment was performed overnight, with DMSO serving as a control.

### 2.5 Preparation and characterization of antibody drug conjugates

Monomethyl auristatin E (MMAE) conjugated 1A3 antibody (1A3-VC-PAB-MMAE) was prepared using a kit (CM11409) from CellMosaic Inc. (Woburn, MA, USA) following the manufacturer’s instructions. Maleimide-activated valine-citrulline p-aminobenzylcarbamate (VC-PAB) MMAE, was coupled directly to the antibody after reduction through alkylation (Sun et al., 2005). The specificity of 1A3 and 1A3-VC-PAB-MMAE to peptides representing human cleaved-AREG was confirmed by immunoblotting. Briefly, transferrin conjugated peptides representing cleaved and uncleaved AREG (200 ng) were electrophoresed on SDS-PAGE gels, transferred to PVDF membrane and probed with 1 μg/ml of 1A3 and 1A3-VC-PAB-MMAE. Horseradish peroxidase-conjugated goat anti-rabbit IgG was used. Antibody binding was detected with enhanced chemiluminescence (ECL; Amersham, Arlington Heights, IL, USA). Furthermore, the characterization and purity of 1A3-VC-PAB-MMAE was analyzed using hydrophobic interaction chromatography (HIC) and size-exclusion chromatography (SEC) (Cell Mosaic).

### 2.6 Binding affinity of 1A3 and 1A3-VC-PAB-MMAE by ELISA

Maxisorp flat bottom 96-well plates (Thermo Fisher Scientific) were coated with 5 μg/ml cleaved-AREG peptide at 4°C overnight. After washing thrice with PBST (PBS with 0.05% Tween20) and blocking for 2 h at RT with 5% non-fat dry milk in PBST, various concentrations of 1A3 and 1A3-VC-PAB-MMAE (0.001, 0.01, 0.1, 1 μg/ml) were added to the wells and incubated for 2 hours at RT. The wells were washed four times with PBST and were incubated at RT with Goat anti-rabbit IgG (H+L) Cross-adsorbed -HRP conjugate secondary Ab (ThermoFisher G-21234) for 1h. One-Step™ Turbo TMB-ELISA Substrate Solution (Thermo Fisher Scientific) was added to the wells for color development, incubated for 15 minutes and the reaction was stopped by adding 2M H_2_SO_4_. The absorbance was measured at 450 nm using VersaMax Microplate Reader (Molecular Devices LLC, San Jose, California, USA).

### 2.7 In vitro antitumor studies

#### 2.7.1 MTT assay

The cytotoxic effect of 1A3 and 1A3-VC-PAB-MMAE antibodies on MCF7 and MCF7-F breast cancer cells were determined using an MTT (3-(4,5-Dimethylthiazol2-yl)-2,5-Diphenyltetrazolium Bromide) assay. The cells were seeded at a density of 5000 cells/well in 96-well plates. Approximately 24 h after cell seeding, the cells were treated with varying doses (1.56, 3.125, 6.25, 12.5, 25, 50, 100 and 200 nM) of MMAE, 1A3 and 1A3-VC-PAB-MMAE. Untreated cells were used as control. After incubating for 72 h, the medium was aspirated, and 50 μL of serum-free media and 50 μL of MTT solution (5 mg/mL solution in PBS) was added into each well; the cells were then incubated for 3 hours at 37°C. The formazan crystals formed were dissolved with 4 mM HCl, 0.1% NP40 in isopropanol. The absorbance was measured at 590 nm using VersaMax Microplate Reader (Molecular Devices LLC, San Jose, California, USA). The assays were performed in triplicate.

#### 2.7.2 Clonogenic assay

MCF7 breast cancer cells were plated in 48-well plates, allowed to attach overnight and treated with 12.5, 25, or 50 nM of 1A3-VC-PAB-MMAE for 72 h. A vehicle treated well served as control. After removing the ADC-containing medium, cells were washed using PBS, trypsinized and plated at a density of 1000 cells/well in 12-well plates. The cells were cultured for 14 days, with fresh medium addition every three days. The colonies were stained with 0.5% crystal violet solution. The number of colonies per well were counted.

### 2.8 In vivo antitumor activity studies

The animal experiments were approved by the Institutional Animal Care and Use Committee (IACUC) of University of Wisconsin La Crosse (La Crosse, Wisconsin, USA). Four-week-old female NOD.Cg-Prkdc^scid^ Il2rg^tm1Wjl^/SzJ mice (NSG; The Jackson Laboratory) were acclimated for 1 week before the experiments. For MCF7 breast cancer xenograft model, the mice were implanted subcutaneously with 0.72-mg 17β-estradiol 90-day release tablets (Innovative Research of America) three days prior to cell injection. The endocrine resistant MCF7-F xenograft model did not require 17β-estradiol pellets. MCF7 and MCF7-F cells were infected with pLenti-CMV-puro-Luc lentiviral luciferase construct (Addgene 17477) to monitor tumor growth. Firefly luciferase expressing MCF7 and MCF7-F cells (2×10^6^ cells in a 1:1 mixture of serum free DMEM and Matrigel) were injected orthotopically into the fourth inguinal mammary fat pad of each mouse. Once the tumors were well established, each cell line cohort was divided into three groups (n= 5). Mean tumor volume at the start of treatment was 111 mm^3^ for the MCF7 cohorts and 119 mm^3^ for the MCF7-F cohorts. The mice were treated by intraperitoneal injection every four days for a total of six doses with unconjugated or MMAE conjugated anti-cleaved-AREG 1A3 antibody (5 mg/kg), or vehicle control (PBS). Tumor volumes were calculated from caliper data by using the formula: Volume (mm^3^) = (Length x Width^2^)/2. Mice were imaged weekly with the In-Vivo FX PRO imaging system (Carestream Molecular Imaging, Woodbridge, CA, USA,) after injecting luciferin (150 mg/kg body weight). Tumor volume and body weight were monitored once a week.

### 2.9 Statistical Analysis

Statistical analyses were performed using Graph Pad Prism 7 (San Diego, CA, USA) software. Comparisons between groups were made using unpaired t-test (two groups) or one-way/two-way ANOVA (three or more groups) with Bonferroni (for intergroup comparisons) or Dunnett’s (when comparing relative to a single control) multiple comparison tests. Spearman’s rank order coefficient correlation was used to analyze full length vs. cleaved AREG IHC results. P values <0.05 were considered as significant and are indicated by asterisks in figures (****, P < 0.0001; ***, P < 0.001; **, P < 0.01; *, P < 0.05).

## Results

### Candidate Antibody Identification

The EGFR-activating signaling domain of Amphiregulin is inducibly released by ADAM17-mediated cleavage and the residual transmembrane stalk is internalized by endocytosis (Fig 1A). We previously reported the N-terminal sequence of this residual transmembrane Amphiregulin fragment (Levano and Kenny, 2012). Reasoning that antibodies selectively recognizing this sequence in its cleaved but not uncleaved conformation might provide useful therapeutic reagents, we used phage display to isolate rabbit VK-linker-VH scFVs with these binding characteristics (Fig 1B). After IgG reformatting of the most favorable hits, we obtained three monoclonal rabbit IgG antibodies that selectively recognized a peptide representing the neo-epitope of the cell membrane-bound ADAM17-cleavage product of human amphiregulin (Fig 1C). We pursued further study with 1A3, the candidate antibody which exhibited the highest binding ratio of cleaved AREG to intact AREG as determined by ELISA.

**Figure 1.**
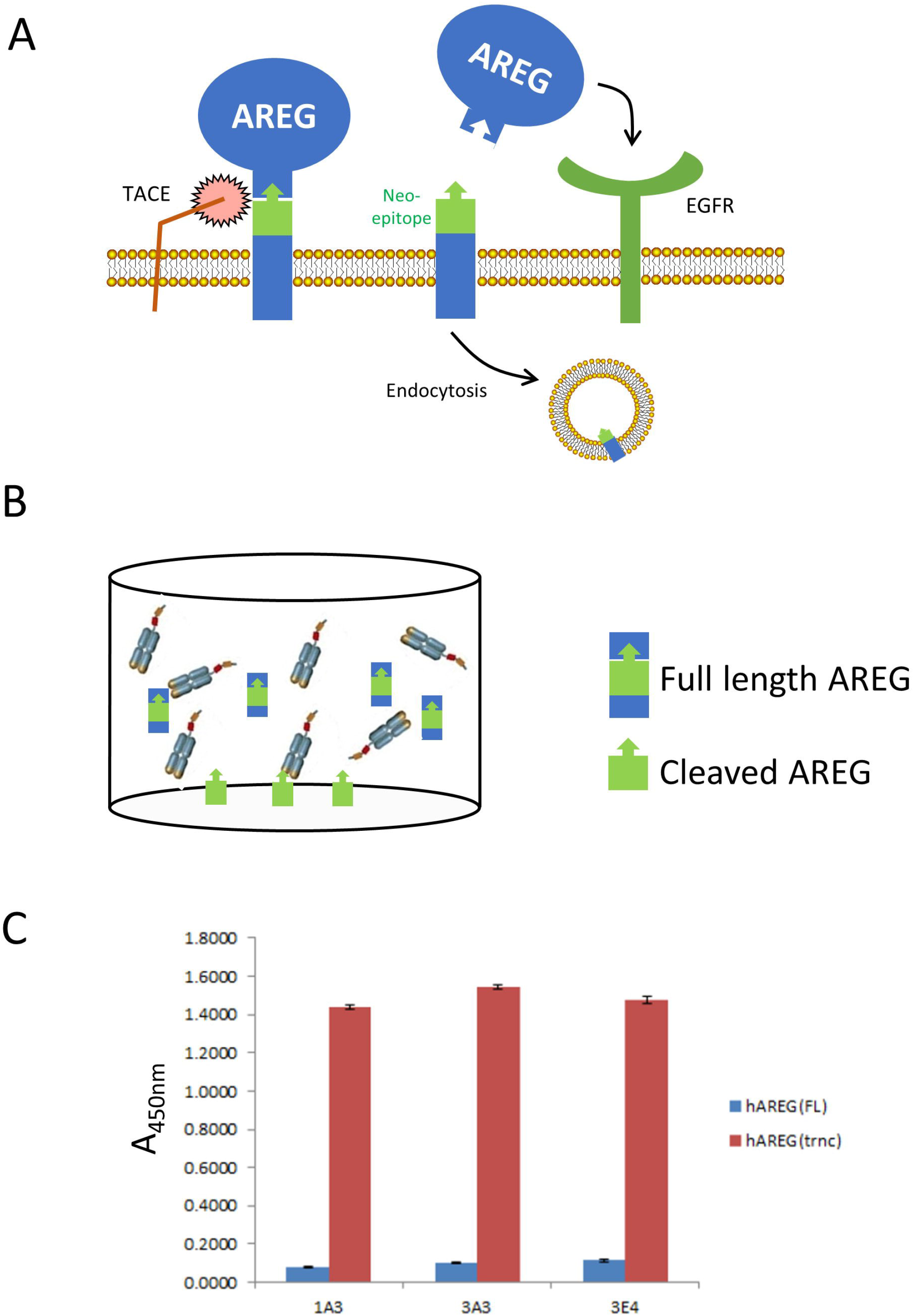
Isolation of antibodies that selectively recognize a peptide representing cleaved Amphiregulin. A. Schematic representation of the Amphiregulin cleavage and EGFR activation process, one consequence of which is the internalization of the small transmembrane fragment that remains cell associated after TACE/ADAM17-mediated cleavage. B. Schematic representation of phage panning strategy using a peptide representing the cleaved Amphiregulin neo-epitope adhered to the well and a peptide representing the non-cleaved Amphiregulin region in the liquid phase. C. ELISA analysis of the three lead candidate rabbit IgG molecules demonstrating their relative affinities for peptides representing the full length and cleaved/truncated Amphiregulin.

### Internalization of 1A3 antibody is dependent on AREG cleavage

Having demonstrated that the 1A3 antibody had the requisite selectivity profile against synthetic peptides (Fig 1C), we then evaluated whether it could recognize endogenous cleaved Amphiregulin in its cellular context and, further, if it could be used to internalize a cargo. Conjugation with pHrodo, a dye that fluoresces in acidic environments, was used to evaluate both cell binding and internalization. We treated MCF7 cells with Hoechst and Lysotracker to visualize the nuclei and lysosomes, respectively, and then exposed them to 1A3-pHrodo. Red fluorescence, indicating presence of 1A3 pHrodo in acidic endosomes was evident after 30 minutes of 1A3-pHrodo treatment and increased in intensity over time (Fig 2A). Colocalization of pHrodo and Lysotracker signals also increased throughout the time course, demonstrating that 1A3 was trafficked to the lysosome after internalization, a subcellular destination necessary for linker cleavage and release of the active drug (Schrama et al., 2006).

**Figure 2.**
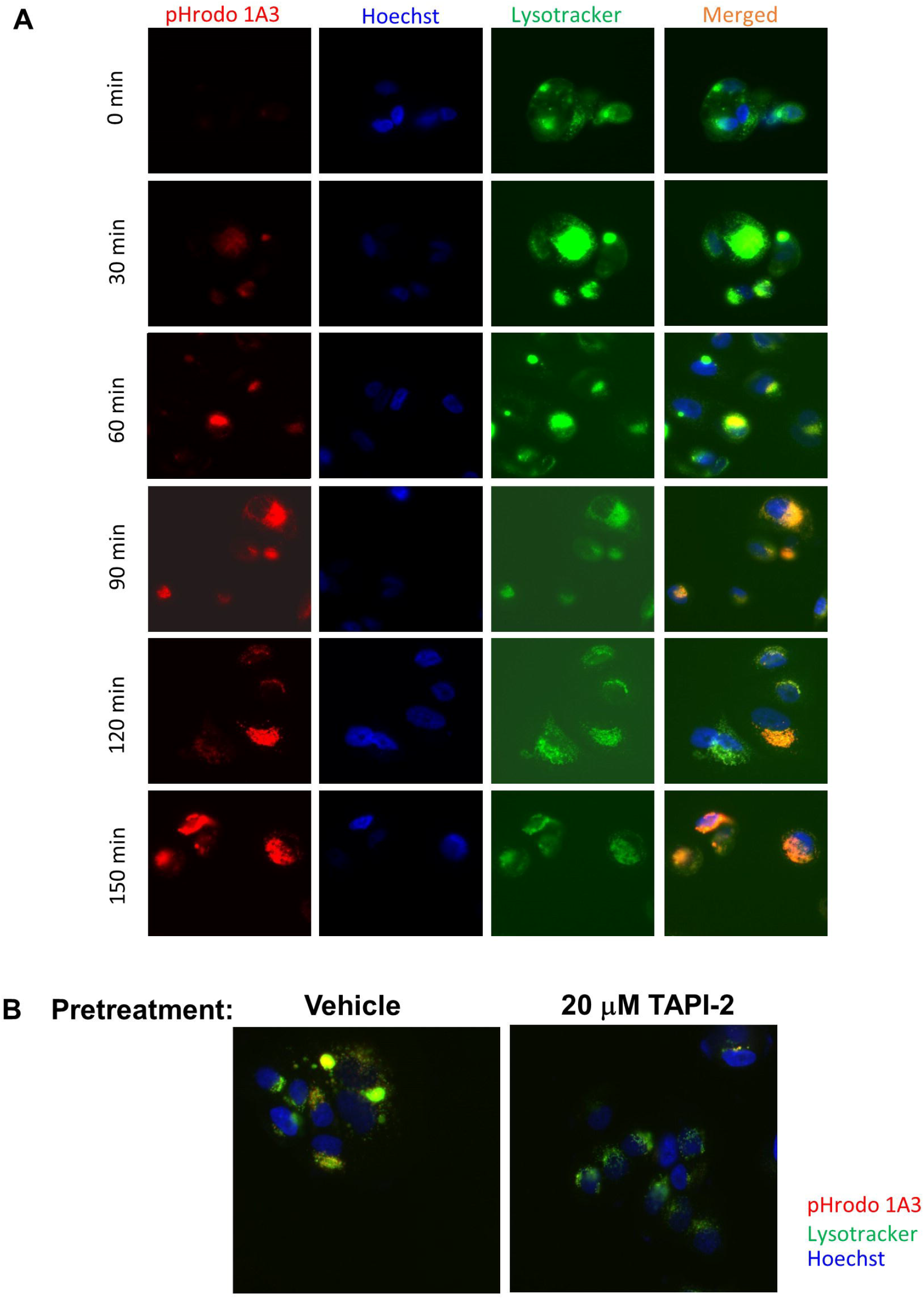
1A3 recognizes the cleaved neo-epitope in its cellular context and can be used to internalize a fluorescent cargo. A. MCF-7 cells labeled with Hoechst 33342 (blue) and Lysotracker (green) were treated with 1A3-pHrodo (red) and monitored with live cell imaging. Note the appearance of yellow in the merged image indicating trafficking of the antibody to lysosomes. B. MCF-7 cells were pretreated with TAPI-2 overnight to prevent AREG cleavage, labeled with Hoechst and Lysotracker, and imaged as in A. Merged images are shown. Comparative absence of red and yellow signal in the TAPI-2 treated cells shows failure to internalize 1A3-pHrodo.

To determine whether ADAM17-mediated proteolytic generation of this neo-epitope was required for 1A3 internalization, we pretreated cells with the ADAM17 inhibitor, TAPI-2. This prevented the development of fluorescence after cells were incubated with 1A3-pHrodo treatment (Fig 2B), from which we conclude that internalization of 1A3-pHrodo requires the generation of the neo-epitope by proteolytic cleavage.

### Retention of epitope selectivity following MMAE conjugation

Using MMAE, we generated an antibody drug conjugate from 1A3 in order to assess the cytotoxicity in vitro and in vivo. To evaluate whether 1A3-MMAE retained the ability to selectively recognize the AREG neo-epitope after conjugation, immunoblots with 1A3 or 1A3-MMAE were performed with AREG peptides representing either an intact ADAM17-cleavage sequence (FL AREG) or the fragment that remains cell-bound after cleavage by ADAM17 (Trunc AREG). Both 1A3 and 1A3-MMAE specifically recognized the truncated AREG and also, as expected, did not react with the peptide representing uncleaved AREG (Fig 3A). A capture ELISA using the cleaved AREG peptide also exhibited no substantial difference in binding between the unconjugated 1A3 and 1A3-MMAE (Fig. 3B). HIC-HPLC of 1A3-MMAE (red) exhibited increased retention time relative to the unconjugated 1A3 (blue), resulting in a calculated average drug antibody ratio (DAR) of 4.4 (Fig 3C). This placed our ADC within the range of several previously investigated ADCs with DARS between 2-6 that was strikes a balance between ADC solubility with an effective degree of payload delivery (Akaiwa et al., 2020; Hamblett et al., 2004; Lyon et al., 2015; Sun et al., 2017).

**Figure 3.**
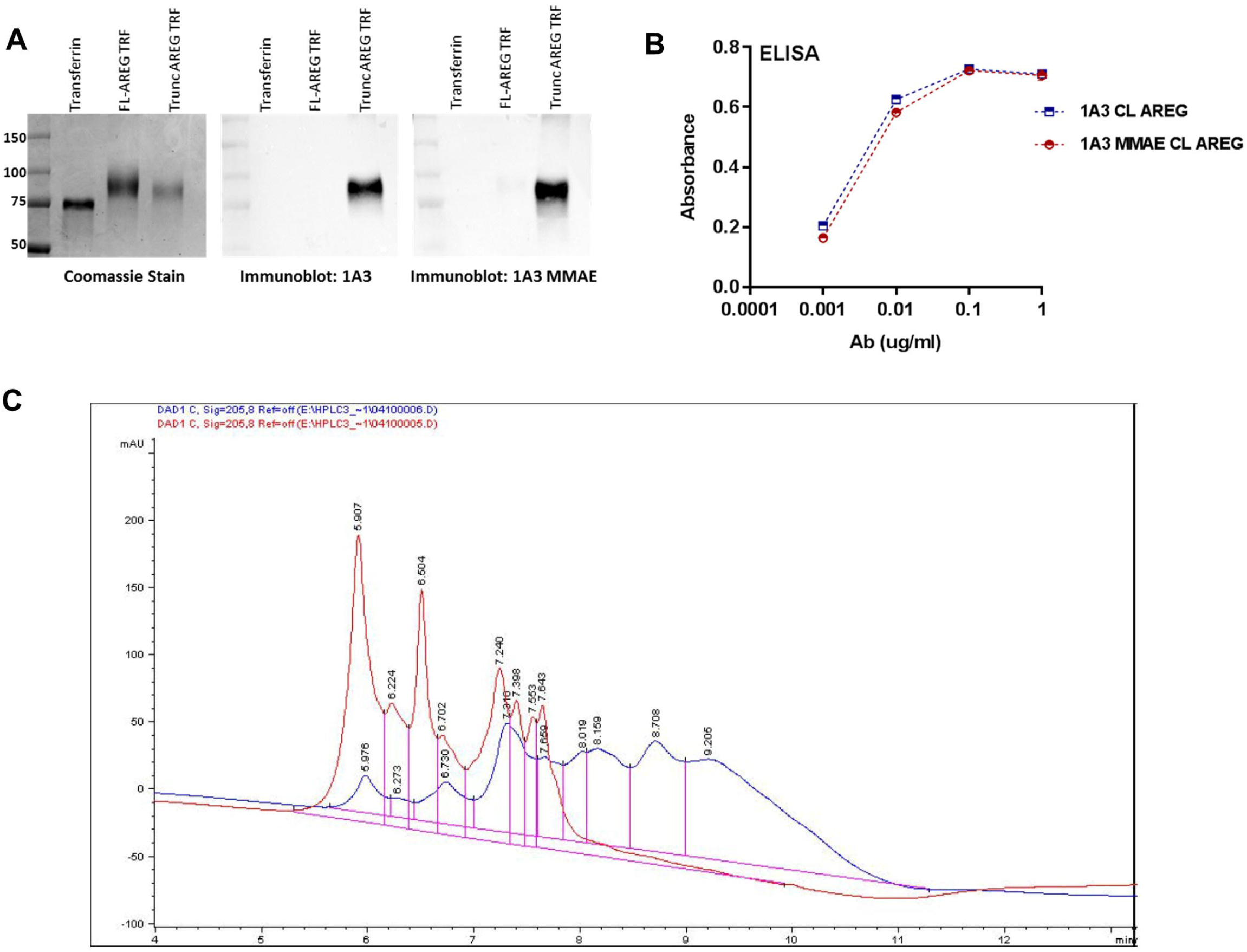
Conjugation of 1A3 with MMAE does not disrupt target recognition. A. SDS-PAGE of transferrin (TRF), and TRF conjugated AREG peptides including the cleavage site (FL-AREG) or a mimic of cleaved AREG (Trunc AREG). Samples were blotted with 1A3 or 1A3-MMAE antibody. B. ELISA comparing binding of 1A3 and 1A3-MMAE to a cleaved AREG peptide C. HIC-HPLC profile of unconjugated 1A3 antibody and the 1A3-MMAE conjugate. The average DAR was determined to be 4.4.

### 1A3-MMAE is cytotoxic in breast cancer cell lines

Treating MCF7 and Fulvestrant-resistant MCF7 (MCF7-F) with 1A3-MMAE resulted in considerable cytotoxicity, as measured by MTT assay (Fig 4A, 4B). Importantly, unconjugated 1A3 did not induce cell death, while free MMAE potently killed the cells. Clonogenic assays with 25 nM or 50 nM 1A3-MMAE also showed a dose dependent effect on survival in both MCF7 and MCF7-F cell lines visually (Fig 4C) and when quantified as number of colonies relative to the control (Fig 4D). As MMAE acts to prevent tubulin polymerization, we evaluated microtubule integrity in cells exposed to 1A3-MMAE. Cells that received free MMAE or 1A3-MMAE exhibited tubulin disruption, whereas control treatments with PBS, unconjugated 1A3, or IgG-MMAE did not (Fig 4E).

**Figure 4.**
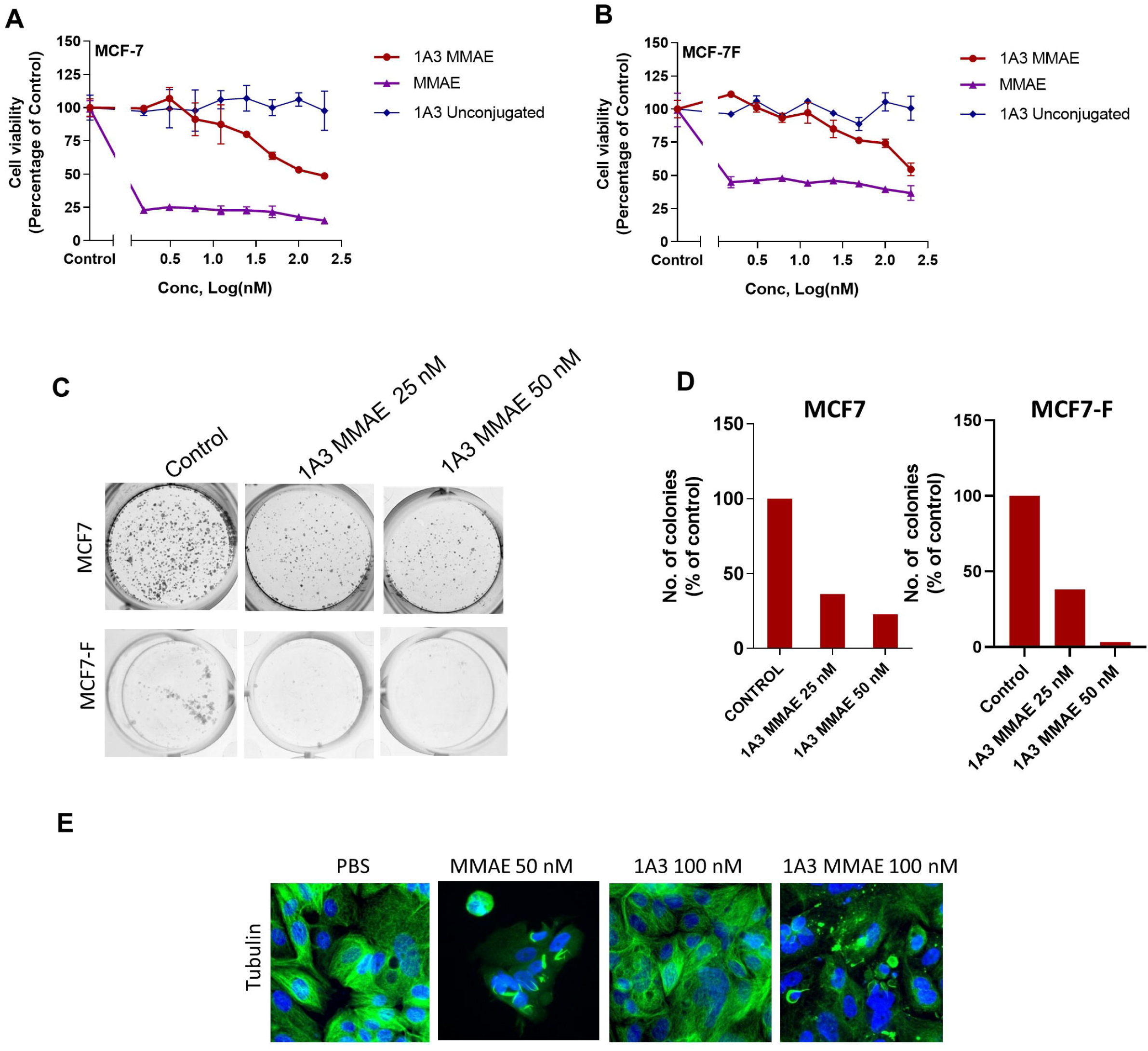
In vitro 1A3-MMAE cytotoxicity. A. Viability of MCF-7 (A) and MCF-7F (B) cells measured by MTT assay after treatment with 1A3-MMAE, 1A3, or MMAE. B. Clonogenic growth of MCF-7 or MCF-7F cells after exposure to 1A3-MMAE and replating. C. Quantification of clonogenic growth assays in C. D. Tubulin staining of MCF-7 cells after 48 hours treatment with MMAE, 1A3, 1A3-MMAE.

### 1A3-MMAE treatment causes regression of breast cancer xenografts

To test *in vivo* efficacy of 1A3-MMAE, human breast cancer cell lines expressing luciferase were injected orthotopically into NSG mice (one tumor in each inguinal mammary gland). Once the average tumor size reached approximately 100-125 mm^3^, mice were randomized to one of three treatment groups: PBS, 1A3 (5 mg/kg) or 1A3-MMAE (5 mg/kg). Mice received intraperitoneal injections every four days for six total injections and were monitored for tumor progression or regression. Caliper measurement of individual MCF7 tumors (Fig 5A) demonstrated strong tumor shrinkage in response to 5 mg/kg 1A3-MMAE. Neither the PBS nor unconjugated 1A3 treated groups showed a disruption in tumor growth with continued, sometimes rapid increases in tumor volume relative to the 1A3-MMAE group.

**Figure 5:**
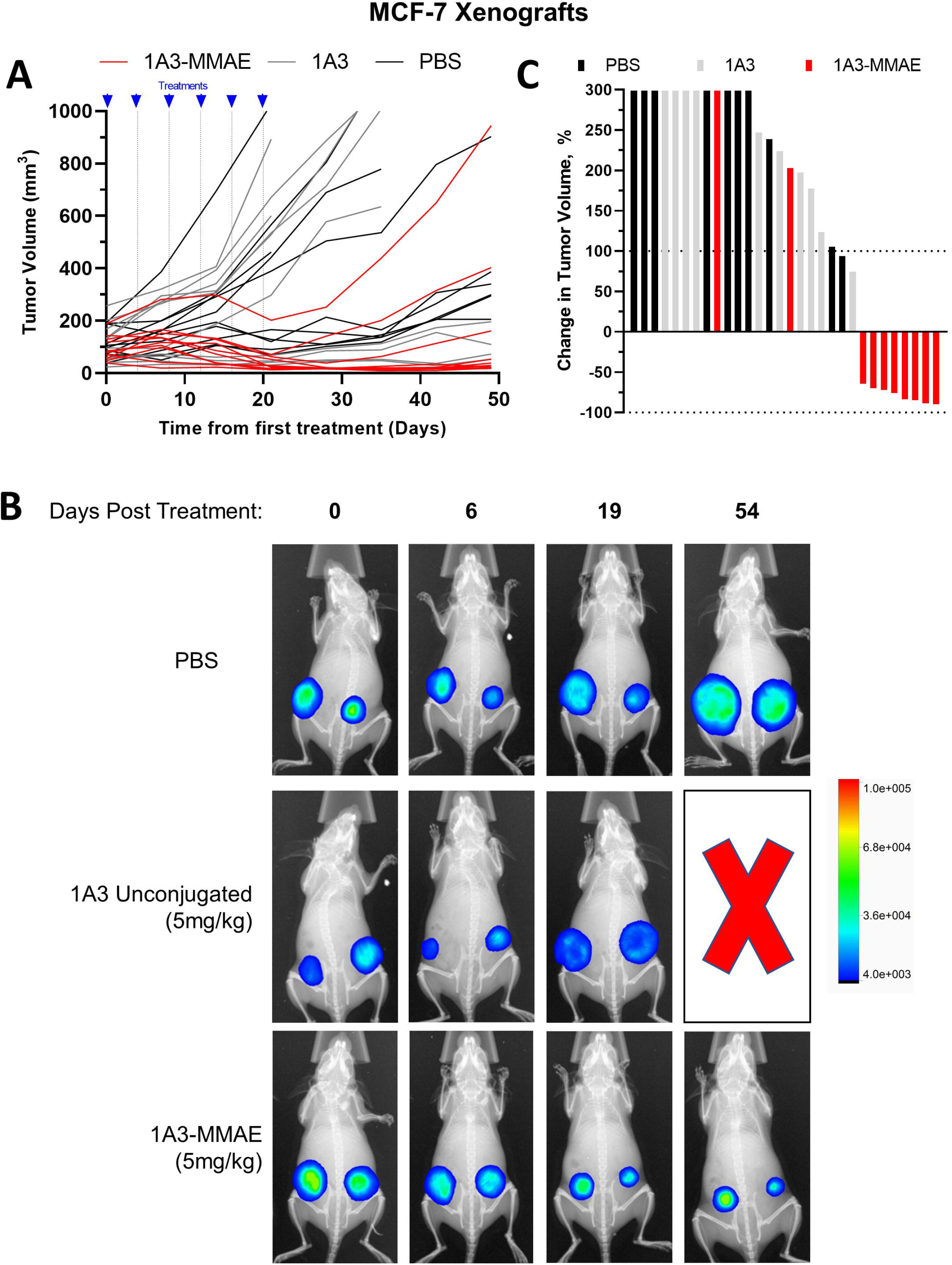
In vivo anti-tumor efficacy in MCF7. A. Caliper measurements of MCF-7-luc orthotopic xenografts after treatment initiation. Each curve represents an individual tumor. Treatment times for the six treatments are indicated. B. Representative *in vivo* luminescence of MCF7 xenograft tumors from A during the course of treatment. A red X indicates an animal euthanized due to tumor burden. The heatmap indicates the intensity of the luminescence signal C. Waterfall plot showing maximum response of the MCF7 xenograft tumors to the treatment course. The Y-axis is truncated at 300%.

Longitudinal luciferase imaging of established xenografts and their response to treatments (Fig 5B) showed changes in tumor size that matched volume trends seen with caliper measurements. One representative animal is shown from each group. The PBS and unconjugated 1A3 groups exhibited tumor growth evident by increased lesion diameter and increasing signal intensity *(color change)*. In contrast, tumors in the 1A3-MMAE treated groups both decreased in both volume and signal. Similar results were seen with MCF7-F xenografts (Figs 6A, 6B). Notably, in the MCF7-F/1A3-MMAE image, one can see the tumor signal decreasing below the limit of detection for imaging, and with continued follow-up, 70 days after the end of dosing, the re-appearance of a lesion.

**Figure 6:**
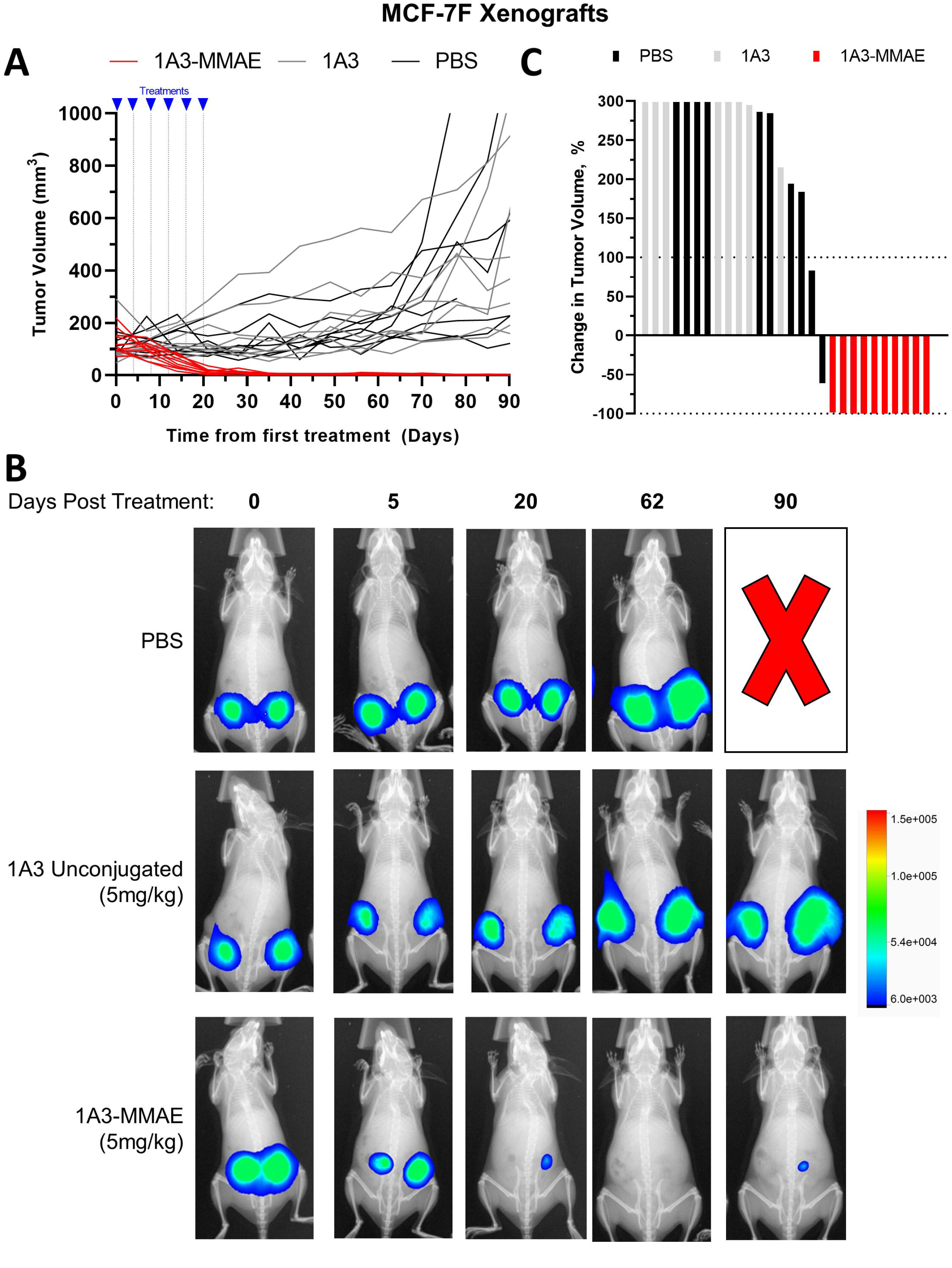
In vivo anti-tumor efficacy in endocrine resistant MCF7-F. A. Caliper measurements of MCF7-F-luc orthotopic xenografts after treatment initiation. Each curve represents an individual tumor. Treatment times for the six treatments are indicated. B. Representative *in vivo* luminescence of MCF7-F xenograft tumors from A during the course of treatment. A red X indicates an animal euthanized due to tumor burden. The heatmap indicates the intensity of the luminescence signal C. Waterfall plot showing maximum response of the MCF7-F xenograft tumors to the treatment course. The Y-axis is truncated at 300%.

To evaluate overall efficacy, we determined the maximum response as defined by the maximum change in tumor volume relative to pre-treatment measurements, depicted as waterfall plots for MCF7 (Fig 5C) and MCF7-F (Fig 6C). Of the MCF7 xenografts, tumor volume increased in 2/10 and decreased in 8/10 dosed with 5 mg/kg 1A3-MMAE. As the 5 mg/kg dose of 1A3-MMAE successfully cleared the tumors to sub-detectable levels in all 10 of the MCF7-F xenografts, we continued follow up in this cohort to determine whether this was curative or if recurrence was possible. After a 204 day follow up period, 10/10 tumors had recurred in mice treated with 5 mg/kg 1A3-MMAE, however only one tumor treated with 1A3-MMAE returned to a size seen prior to treatment, with an average time to recurrence of 73 days after initial tumor clearance determined by luciferase imaging.

### Evaluation of 1A3 utility as a companion diagnostic

We performed immunohistochemistry on a breast tumor tissue micro array to obtain an estimate of the prevalence of the target in breast cancer and to determine whether 1A3 staining had potential for use as a treatment selection biomarker. In general, samples with high AREG expression usually exhibited high cleaved AREG intensity (Fig 7A, 7B). This indicates that Amphiregulin-high tumors (1) generally actively cleave Amphiregulin, (2) that the recycling kinetics in tumors are not so rapid as to preclude detection of the epitope by these antibodies and (3) that these antibodies may have additional utility as companion diagnostics. Among ER-positive (n = 88) and ER-negative (n = 50) tumors evaluated, the proportions of tumors exhibiting medium/high intensity staining with 1A3 were essentially equivalent (69.3% v 70%, respectively). While we have previously reported that Amphiregulin tends to be more highly expressed in ER-positive breast tumors (Peterson et al., 2015), there is clearly a substantial fraction of ER-negative tumors which express and cleave Amphiregulin in sufficient quantities to be considered therapeutic candidates.

**Figure 7.**
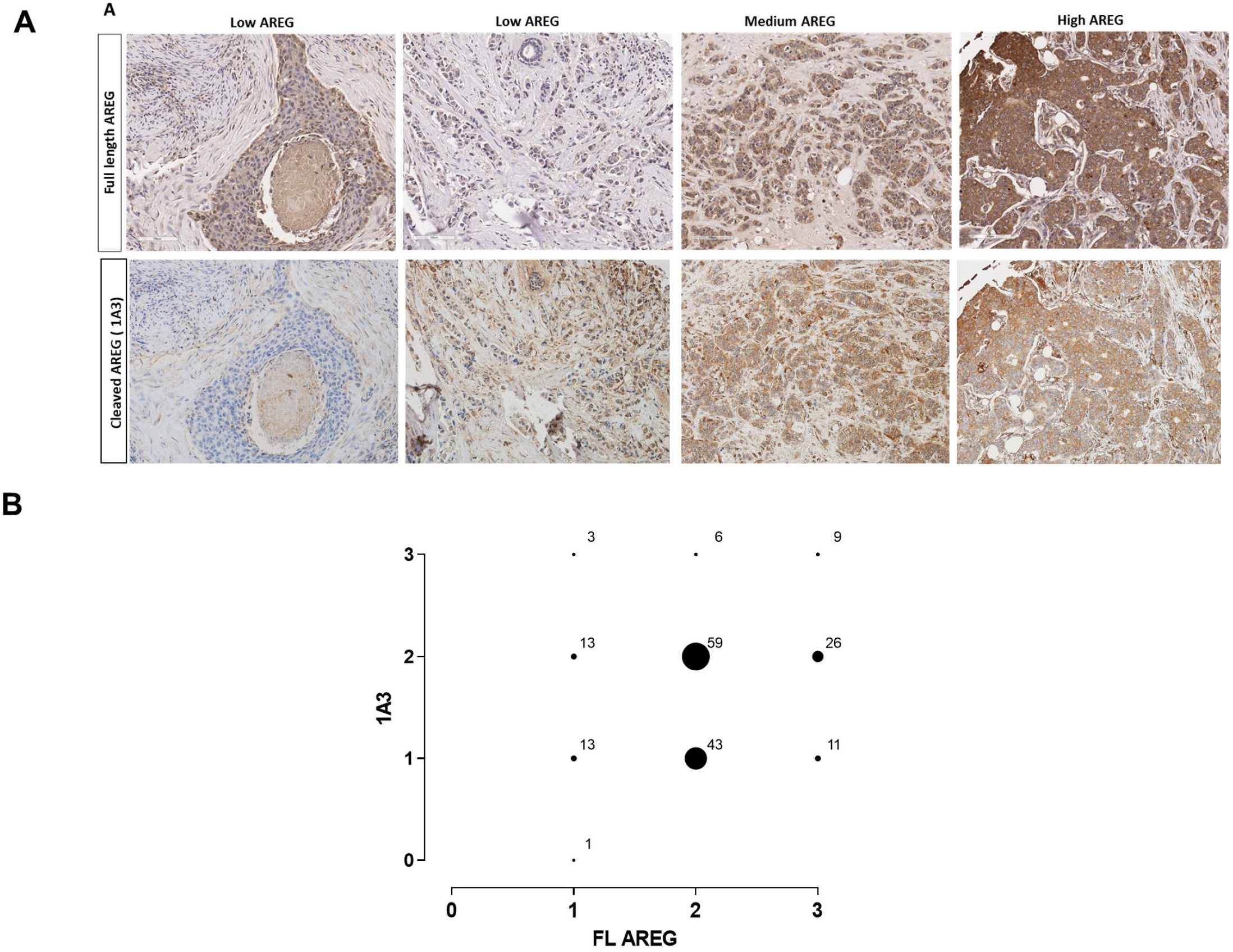
Evaluation of 1A3 antibody utility as a companion diagnostic. A. Representative examples of immunohistochemical staining intensities observed in a human breast tissue array using an antibody to full-length AREG (top row), or 1A3 targeting cleaved AREG (bottom row). B. Cross-comparison of intensity scores for full-length (FL) and cleaved AREG (1A3) from the breast tissue array.

## Discussion

In this study, we establish that the cell-associated transmembrane stalk of Amphiregulin that is generated by ADAM17-mediated cleavage is a viable therapeutic target in breast cancer. We identified three monoclonal antibodies that very selectively recognized this neo-epitope after cleavage, and which did not interact with full-length Amphiregulin. Choosing an antibody, 1A3, which had marginally lower background signal against the uncleaved AREG peptide than the other two candidates (Fig 1C), we performed a series of in vitro and in vivo experiments to evaluate therapeutic potential. By conjugation with a pH-sensitive dye (Fig 2A), we showed that 1A3 is efficiently internalized and trafficked to the lysosome (a cellular endpoint that is a necessary pre-requisite for ADC efficacy) and that this internalization is dependent on cleaved Amphiregulin (Fig 2B). We showed that conjugating with MMAE did not alter target recognition or selectivity (Fig 3A, 3B). This antibody drug conjugate, 1A3-MMAE, efficiently disrupted the tubulin cytoskeleton of breast cancer cells (Fig 4E) resulting in cell death in vitro (Fig 4A-D). In vivo, 1A3-MMAE caused substantial regression of tumors formed by xenografted human breast cancer cell lines (Fig 5, Fig 6). In all experiments, the naked antibody was ineffective, demonstrating that it is the target mediated MMAE internalization that underlines the therapeutic effect.

Identifying an appropriate target antigen is critical for efficacy and tolerability of ADC dosing in patients; ideally one would choose a tumor-specific antigen rather than a tumor-associated antigen to decrease the likelihood of on-target, off-tumor effects. Here, by demonstrating the feasibility of designing an ADC against a transient protein cleavage product, our study suggests that developing ADCs against targets that display different epitopes in some manner dependent on their state of activation/processing may generally provide even more selective ways to target relevant cell populations in vivo. Human and mouse Amphiregulin have some amino acid sequence differences in the region of the neo-epitope making it unlikely that our antibody recognizes endogenous mouse Amphiregulin. Accordingly, although we did not observe weight loss, lethargy or breathing difficulty not attributed to tumor burden in any treated animals, we cannot draw any inference from these data about the potential for toxicity in humans.

Our two series of in vivo experiments differed in one important respect. MCF7 growth required supplementation with slow-release estrogen pellets while the endocrine resistant MCF7-F cell line grows independently of estrogen and these animals were not supplemented. Although both groups of tumors responded well to 1A3-MMAE, the MCF7-F tumors experienced more pronounced responses. Whether this can be explained by the difference in hormonal milieu, a greater dependence on Amphiregulin in this endocrine resistant cell line or some other reason is unclear. Nevertheless, the strong response observed in the endocrine resistant line suggests that these ADCs may warrant evaluation in this patient population for whom there are fewer effective treatments available.

Another anti-AREG therapeutic antibody strategy was recently reported (Lindzen et al., 2021) in which antibodies recognizing the shed Amphiregulin signaling domain can be used to stoichiometrically bind and sequester Amphiregulin released by ovarian cancer cells, thereby blocking autocrine signaling. The approach outlined in our study has the potential advantage of directly killing the Amphiregulin releasing cells however, given that off-target toxicities with an ADC (if they occur) are likely to be more severe, this study offers a useful alternative strategy for targeting Amphiregulin, as well as establishing ovarian cancer as another malignancy in which this EGFR ligand may play an important role.

An appropriate companion diagnostic is critical to the successful development of a targeted therapeutic. We have shown that the 1A3 antibody efficiently detects cleaved Amphiregulin in formalin fixed paraffin embedded breast cancer specimens (Fig 7). Importantly, this series demonstrated that while cleaved AREG is quite commonly expressed in ER+ tumors, it is also found in ER-tumors. Accordingly, while our experimental focus has been on ER+/AREG+ tumors, it is reasonable to speculate that the potential utility may extend to ER-/AREG+ tumors or, indeed, to AREG-high tumors in other tissue types (Busser et al., 2011).

In conclusion, this study demonstrates the feasibility of generating selective antibody-drug conjugates against transient neo-epitopes generated by post-translational processes like proteolytic cleavage. We present a novel anti-Amphiregulin antibody-drug conjugate with potential utility in breast cancer treatment.

## Acknowledgments

This study was supported by the Department of Defense Breast Cancer Research Program (W81XWH-14-1-0294 to PK) and the Gundersen Medical Foundation. KL was supported by the Norman L. Gillette, Jr. Breast Cancer Research Fellowship. PK holds the Dr. Jon and Betty Kabara Endowed Chair of Precision Oncology.

## Notes

### Competing Interest Statement

The authors have declared no competing interest.

